# Molecular and functional variation in iPSC-derived sensory neurons

**DOI:** 10.1101/095943

**Authors:** Jeremy Schwartzentruber, Stefanie Foskolou, Helena Kilpinen, Julia Rodrigues, Kaur Alasoo, Andrew J Knights, Minal Patel, Angela Goncalves, Rita Ferreira, Caroline Louise Benn, Anna Wilbrey, Magda Bictash, Emma Impey, Lishuang Cao, Sergio Lainez, Alexandre Julien Loucif, Paul John Whiting, HIPSCI Consortium (www.hipsci.org), Alex Gutteridge, Daniel J Gaffney

**Author notes:** Corresponding authors: Jeremy Schwartzentruber, Alex Gutteridge, Daniel Gaffney.

## Abstract

Induced pluripotent stem cells (iPSCs), and cells derived from them, have become key tools to model biological processes and disease mechanisms, particularly in cell types such as neurons that are difficult to access from living donors. Here, we present the first map of regulatory variants in an iPSC-derived cell type. To investigate genetic contributions to human sensory function, we performed 123 differentiations of iPSCs from 103 unique donors to a sensory neuronal fate, and measured gene expression, chromatin accessibility, and neuronal excitability. Compared with primary dorsal root ganglion, where sensory nerves collect near the spinal cord, gene expression was more variable across iPSC-derived neuronal cultures, particularly in genes related to differentiation and nervous system development. Single cell RNA-sequencing revealed that although the majority of cells are neuronal and express the expected marker genes, a substantial fraction have a fibroblast-like expression profile. By applying an allele-specific method we identify 3,778 quantitative trait loci influencing gene expression, 6,318 for chromatin accessibility, and 2,097 for RNA splicing at FDR 10%. A number of these overlap with common disease associations, and suggest candidate causal variants and target genes. These include known causal variants at *SNCA* for Parkinson’s disease and *TNFRSF1A* for multiple sclerosis, as well as new candidates for migraine, Parkinson’s disease, and schizophrenia.

## Introduction

Cellular disease models are critical for understanding the molecular mechanisms of disease and for the development of novel therapeutics. Advancements in induced pluripotent stem cell (iPSC) technology are enabling the development of these models in many human cell types. Initial uses of iPSCs for disease modeling have focused mostly on highly penetrant, rare coding variants with large phenotypic effects, such as mutations of IKBKAP in familial dysautonomia (Lee et al. 2009), mutations of SCN9A in chronic pain due to erythromelalgia (Cao et al. 2016), and others (Itzhaki et al. 2011; Liu et al. 2011; Wainger et al. 2014). However, there is growing interest in using iPSCs to model the effects of common genetic variants that drive complex disease. Regulatory genetic variants have been identified for gene expression (The GTEx Consortium et al. 2015), chromatin accessibility (Degner et al. 2012), and transcription factor binding (Ding et al. 2014; Tehranchi et al. 2016) in both primary tissues and immortalized cell lines. However, for many tissues, pure cultures of individual cell types are either unavailable or are too scarce to enable a variety of molecular assays to be performed. iPSC-derived cells are a renewable source of cells which can be genetically manipulated to investigate causal genetic effects.

A key question is to what extent variability in directed differentiation, whether due to stochastic factors or cell line differentiation capacity, is a barrier to studying the effects of common disease-associated variants in iPSC-derived cells (Rouhani et al. 2014; Kajiwara et al. 2012). Common traits and diseases are influenced by hundreds of genetic variants, each typically of modest effect size (Manolio et al. 2009). Observing the molecular phenotypes associated with a disease depends on having the relevant cell type, and on the genetic effects of interest being observable in those cells. Because cultured cells are imperfect models of primary tissues, it is critical to understand which common disease-associated genetic variants also alter cell phenotypes in iPSC-derived systems.

Whereas pain sensation has largely been studied in rodent models, the development of efficient protocols to differentiate iPSCs into nociceptive (pain-sensing) neurons (Young et al. 2014) provides the opportunity to model common genetic effects on human sensory neuron function, which may underlie individual differences in pain sensitivity and chronic pain. Chronic pain is a common complex disease with moderate heritability of ~38% (McIntosh et al. 2016), which is a significant cause of disability globally (Murray et al. 2015). The relevant human tissues for studying pain are the peripheral sensory nerve fibers that innervate the skin and other organs and which are brought together at the dorsal root ganglia (DRG) before synapsing with the spinal cord around the dorsal horn. Because obtaining such tissue is only possible post-mortem, iPSCderived models will be useful tools in revealing pain biology in humans.

Here, we present the first large-scale study of common genetic effects in a cell type differentiated from human stem cells, iPSC-derived sensory neurons (IPSDSNs). Using single-cell RNA sequencing we characterize the heterogeneity in individual cells produced by directed differentiation. We compare variability in gene expression across IPSDSN samples with that seen in primary DRG and in other tissues from the genotype-tissue expression project (GTEx) (The GTEx Consortium et al. 2015). We identify quantitative trait loci (QTLs) where genetic variants influence gene expression, RNA splicing, and chromatin accessibility in these cells and identify a number of overlaps between molecular QTLs and common disease associations, including Parkinson’s disease, migraine, multiple sclerosis, and schizophrenia. In generating this gene regulatory map we establish effective techniques for using IPSDSN cells to model molecular phenotypes relevant to common diseases.

## Results

### Sensory neuron differentiation and characterization

We obtained 103 IPS cell lines derived from unrelated apparently healthy individuals by the HIPSCI resource (Kilpinen et al. 2016), and followed an established small molecule protocol (Young et al. 2014) to differentiate these into sensory neurons of a nociceptor phenotype. We performed a total of 123 differentiations; 16 of these were done with an early version of the protocol (P1) which was subsequently refined (P2) to yield a higher proportion of neuronal cells in the final cultures. Gene expression was measured by RNA-sequencing for all samples; ATAC-sequencing was done to measure accessible chromatin for 31 samples; and neuron electrophysiology was measured by patch clamp and pharmacological modulation for between 31 and 55 samples (Figure 1a). One RNA-seq sample failed sequencing, and four others were outliers based on principal components analysis and were excluded (Supplementary Figure 1). This left a set of 119 differentiations with gene expression data from 100 unique IPS donors for further analysis.

**Figure 1.**
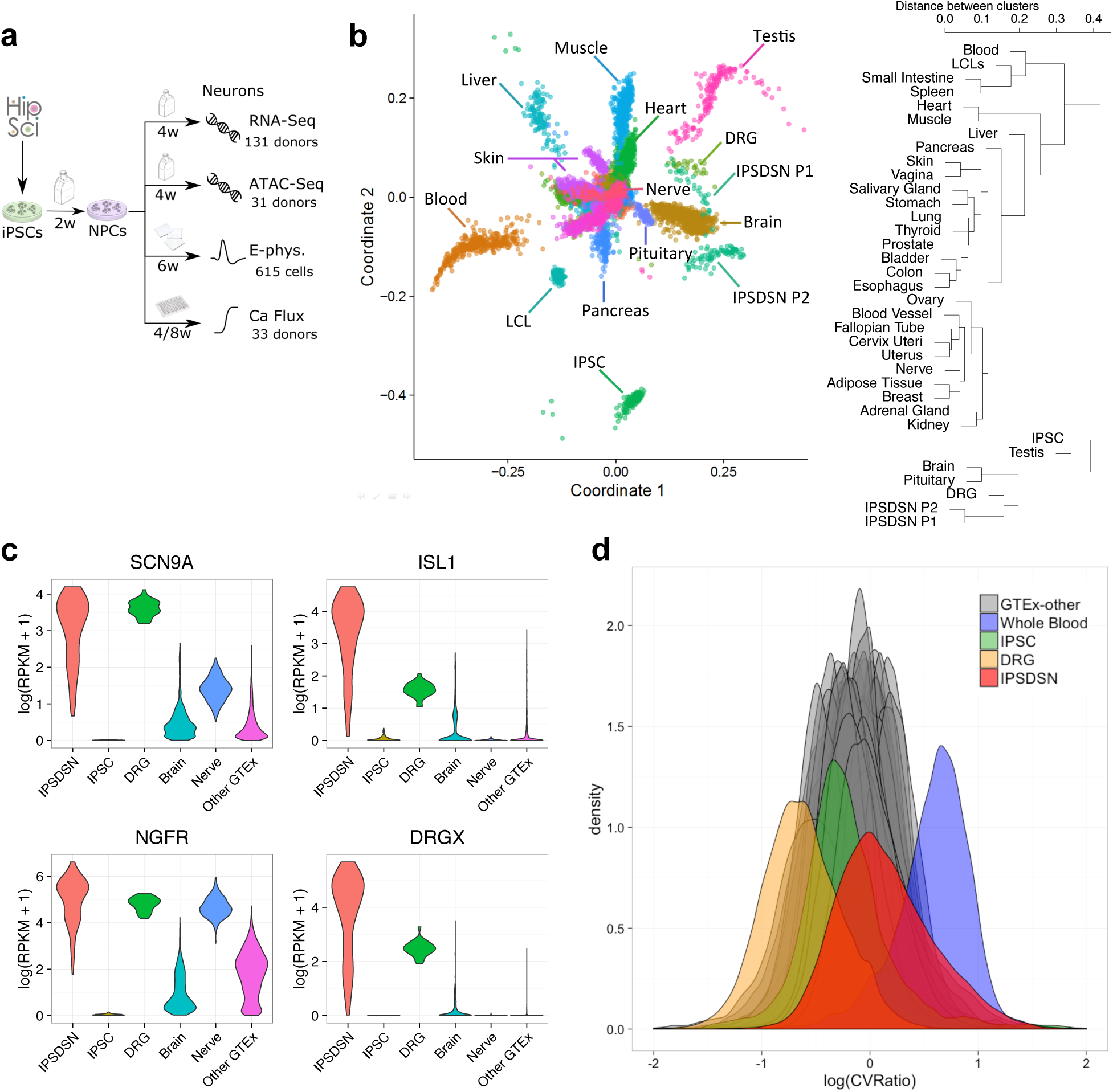
Characterization of molecular phenotypes in IPS-derived sensory neurons. **(a)** Schematic of IPSDSN differentiation and assays, **(b)** Sample and tissue similarity based on gene expression in IPSDSN, iPSC, DRG, and GTEx tissues. Left: Multidimensional scaling, Right: hierarchical clustering, **(c)** Distribution of expression levels for selected sensory neuronal marker genes in IPSDSN, DRG, GTEx tibial nerve, GTEx brain, and all other GTEx tissues, **(d)** Density plot of CVRatio across all genes, separately for each GTEx tissue, IPSDSN samples (n=106, P2 protocol only), iPSC (n=200), and DRG (n=28). CVRatio is the ratio of a gene’s coefficient of variation (CV) across samples in a tissue to the mean CV for that gene in all other tissues.

We first examined the reproducibility of genome-wide gene expression by considering correlations between: replicates of the RNA extraction process within a single differentiation (n=7); replicates of the differentiation process within a single line (n=6 donors, 3 replicates each); and replicates of differentiation across lines (n=94). RNA extraction replicates were highly repeatable (spearman ρ of 0.97 - 0.98). Differentiation replicates within a donor cell line were more variable (median ρ=0.96, range 0.93 - 0.98), but were more highly correlated than differentiations across donors (median ρ=0.93, range 0.80 - 0.98) (Supplementary Figure 2). The variability in gene expression we observed between donor lines could arise from a number of sources, including donor genetic background, effects of clonal selection that occurred during establishment of the line and effects of the cell culture environment during and post reprogramming.

To put the IPSDSN transcriptome in context, we clustered our gene expression data with 200 iPSC samples from many of the same donors, as well as 28 DRG tissue samples from different donors, and 44 primary tissues from the GTEx project (Mele et al. 2015) (Figure 1b). Gene expression was quantified as RNA-seq fragments per kilobase per million reads (FPKM). In hierarchical clustering, IPSDSN samples clustered nearest to DRG, followed by brain samples from GTEx, regardless of clustering method. Marker genes specific to sensory neurons and nociceptors were expressed (FPKM > 1) in nearly all samples (Figure 1c). However, we observed a high degree of heterogeneity in the level of expression of these genes, sometimes over two orders of magnitude (Figure 1c). This contrasted with DRG, where expression of marker genes varied by 3- to 5-fold across samples, despite the fact that a cell culture system is theoretically more pure in cell type composition than a tissue such as DRG. These observations echo difficulties reported elsewhere in using IPS-derived cells to characterize gene expression differences between samples (Soldner et al. 2016; Raghavan et al. 2016).

We examined between-sample variability in global gene expression by computing the coefficient of variation (CV; standard deviation divided by the mean) for each expressed gene (FPKM > 1) among IPSDSN samples (P2 protocol only), and compared this with the same metric for GTEx tissues, DRG, and iPSCs (Figure 1d; Supplementary Figure 3). Whole blood was the most variable tissue, with most genes showing more sample-to-sample variability than the same genes in other tissues, whereas DRG was the least variable tissue. Most genes in IPSDSNs were not highly variable, with the overall distribution of gene expression variability falling within the range of GTEx tissues. However, the median CV of gene expression in IPSDSNs (0.38) was nearly double that in DRG (0.20), indicating that most genes in IPSDSNs have greater sample to sample variability in expression than the primary tissue they are intended to model. Among the technical factors that might explain these differences we saw no effect of sample size or sample RNA integrity on gene expression variability (Supplementary Figures 4,5).

In addition to generally increased expression variability relative to DRGs, we observed a long tail of genes with higher variability in IPSDNs than in nearly all other tissues (Supplementary Table 1). GO enrichment of the top 1000 highly variable genes (CV > 1.28) revealed enrichment for processes relating to cell differentiation and development (Supplementary Table 2), including sensory organ development (4.3x enrichment, p=2×10^−30^), positive regulation of cell differentiation (3.0x enrichment, p=3×10^−24^), and central nervous system development (2.9x enrichment, p=2×10^−22^), although the most highly enriched category was extracellular matrix organization (6.8x enrichment, p=2×10^-54^). Developmental processes were not enriched among the most highly variable genes in DRG, iPSCs, or other GTEx tissues. The iPSCs from which our neurons were derived had gene expression variability within the range seen in GTEx tissues, without a long tail of highly variable genes.

### Single-cell RNA-seq

In previous work we showed that not all individual cells express neuronal marker genes after differentiation (Young et al. 2014). Samples also appeared to differ visually in the fraction of cells with a neuronal morphology (Figure 2b). To further characterize this heterogeneity, we used the Fluidigm C1 with Illumina NextSeq500 to sequence 186 cells from a single IPSDSN sample. After excluding 9 cells expressing fewer than 20% of the ~56,000 quantified genes and noncoding RNAs, we used SC3 (Kiselev et al. 2016) to cluster the remaining 177 cells based on global gene expression. SC3 determines a “consensus” clustering solution by evaluating a range of data transformations and parameter combinations, followed by k-means clustering. The data were best explained by two clusters (Fig 2a and Supplementary Figure 6), with 63% of cells forming a tight cluster expressing sensory-neuronal genes (e.g. *SCN9A, CHRNB2),* and the remaining 37% of cells forming a looser cluster expressing genes typical of a fibroblastic cell type (e.g. *MSN, VIM*). The fibroblast-like cells had ~2.3-fold more RNA-seq reads than sensory-neuronal cells (Supplementary Figure 7), yet expressed fewer genes (>0 fragments overlapping transcript, median 9938 vs. 12750). Although this may indicate greater RNA content in the fibroblast-like cells, it could also reflect differential efficiency in lysing and capturing RNA from these cells relative to the sensory-neuronal cells.

**Figure 2.**
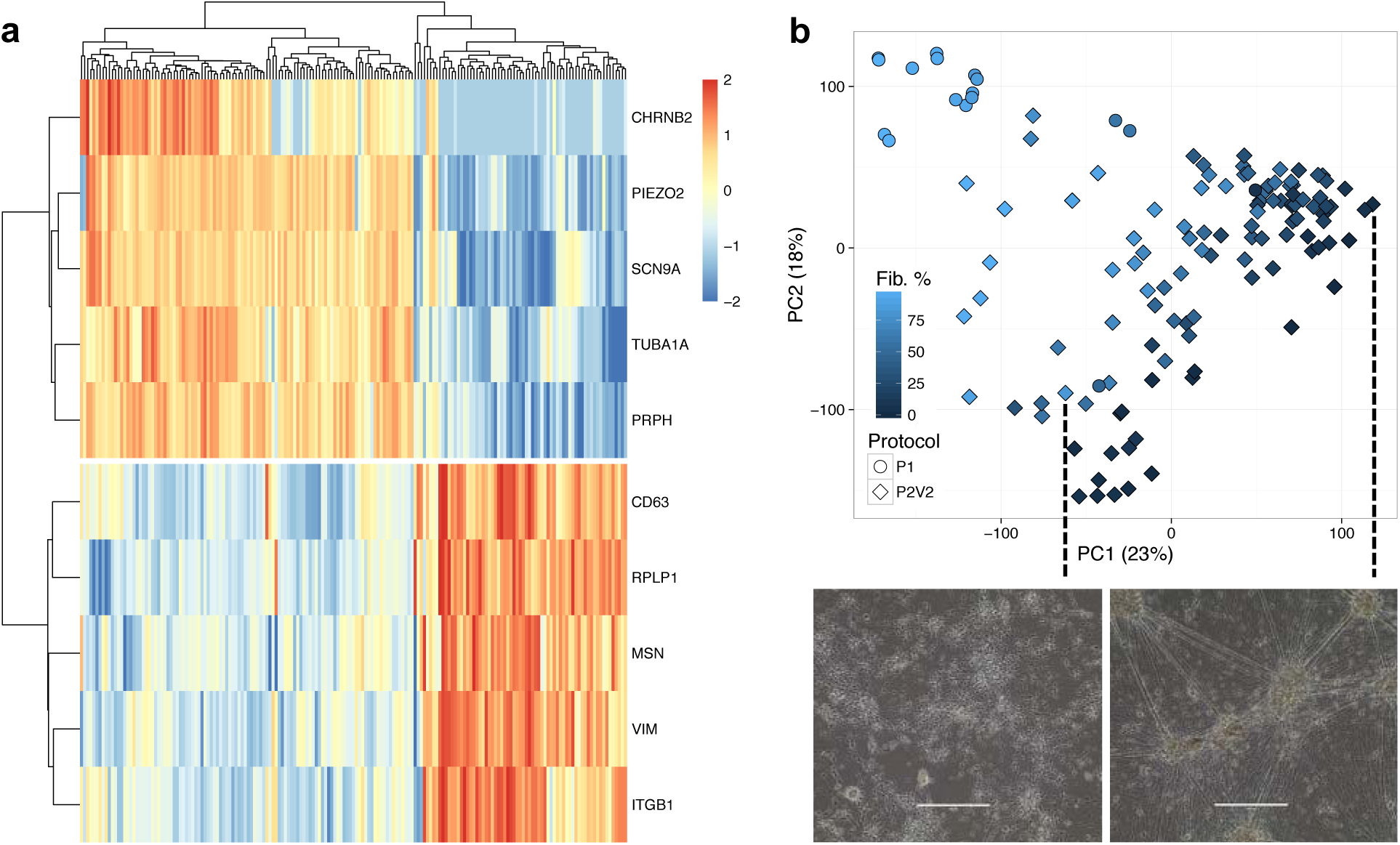
Single-cell sequencing of IPSDSN cells. **(a)** Most cells express markers of sensory neurons, but a significant fraction have fibroblast-like gene expression. **(b)** A PCA plot of RNA-seq samples shows that PC1 separates samples with high fibroblast-like cell content (light blue) from more pure samples with low fibroblast-like cell content (dark blue), based on Cibersort estimates.

The two cell types also separate cleanly in a principal components plot (Supplementary Figure 8), indicating that the cells do not fall on a smooth gradient from more neuronal to less, but rather have differentiated to distinct cell states. Comparing quantile-normalized gene expression from each single cell cluster to other tissues showed that the neuronal cluster is most similar to DRG (Spearman’s ρ=0.751), followed by GTEx brain (mean ρ=0.701) (Supplementary Figure 9). In contrast, the fibroblast-like cluster is most similar to GTEx transformed fibroblasts (ρ=0.768), followed closely by DRG (ρ=0.759). The similarity of the these cells to GTEx fibroblasts could indicate an overall similarity of adherent cultured cells, although the neuronal single cell cluster had lower similarity to GTEx fibroblasts (ρ=0.671) than to a number of other tissues. We proceeded on the basis that single IPSDSN cells can be categorized as either neuronal or fibroblast-like, and that the fraction of neuronal cells may differ between differentiations. We used Cibersort (Newman et al. 2015) to estimate the fraction of RNA from neuronal cells in our bulk RNA-seq samples, using the single cell gene expression counts with their 2-cluster labels as signatures of neuronal vs. fibroblast-like expression. The estimated neuronal content showed a strong correlation with the first principal component of gene expression, and this corresponded well with a visual assessment of neuronal content from microscopy images (Figure 2b, Supplementary Figure 10).

We note that although a large majority of samples appeared by microscopy to have high neuronal content, Cibersort estimated relatively high fibroblast-like content for many samples (median 43%), and as high as 100% for some samples that visually appeared neuronal. A factor contributing to this may be that our scRNA-seq sample was matured for 8 weeks, whereas our bulk RNA-seq samples were matured for 4 weeks. While previous work showed only minor changes in gene expression between 4 and 8 weeks maturation (Young et al. 2014), this difference in maturity means that our single cell reference profiles do not perfectly represent cells in our bulk samples. IPSDSN samples estimated to have high fibroblast content still show greater similarity in genome-wide gene expression with DRG than with any GTEx tissue, including fibroblast cell lines (Supplementary Figures 11, 12). We thus take relative differences between samples in estimated fibroblast-like content as informative, but treat the absolute estimates as uncertain.

### IPSDSN gene expression correlates with functional measurements

We used Ca^2+^ flux measurements on a subset of differentiated cultures (n=31) to confirm that the cells consistently responded to veratridine (a sodium ion channel agonist) and tetrodotoxin (a selective sodium ion channel antagonist), as expected (Supplementary Figure 14). We also performed patch-clamp electrophysiology recordings for 616 individual neurons from 31 donors, with a median of 21 cells measured perline (examples in Supplementary Figures 15,16). For each neuron we measured the resting membrane potential, capacitance and rheobase. The rheobase is the minimum current input that will cause an individual neuron to fire an action potential, and we used this as a measure of the overall membrane excitability. The distribution of rheobases was comparable to those obtained from primary DRG cells, but also showed differences between donors (Figure 3a).

**Figure 3.**
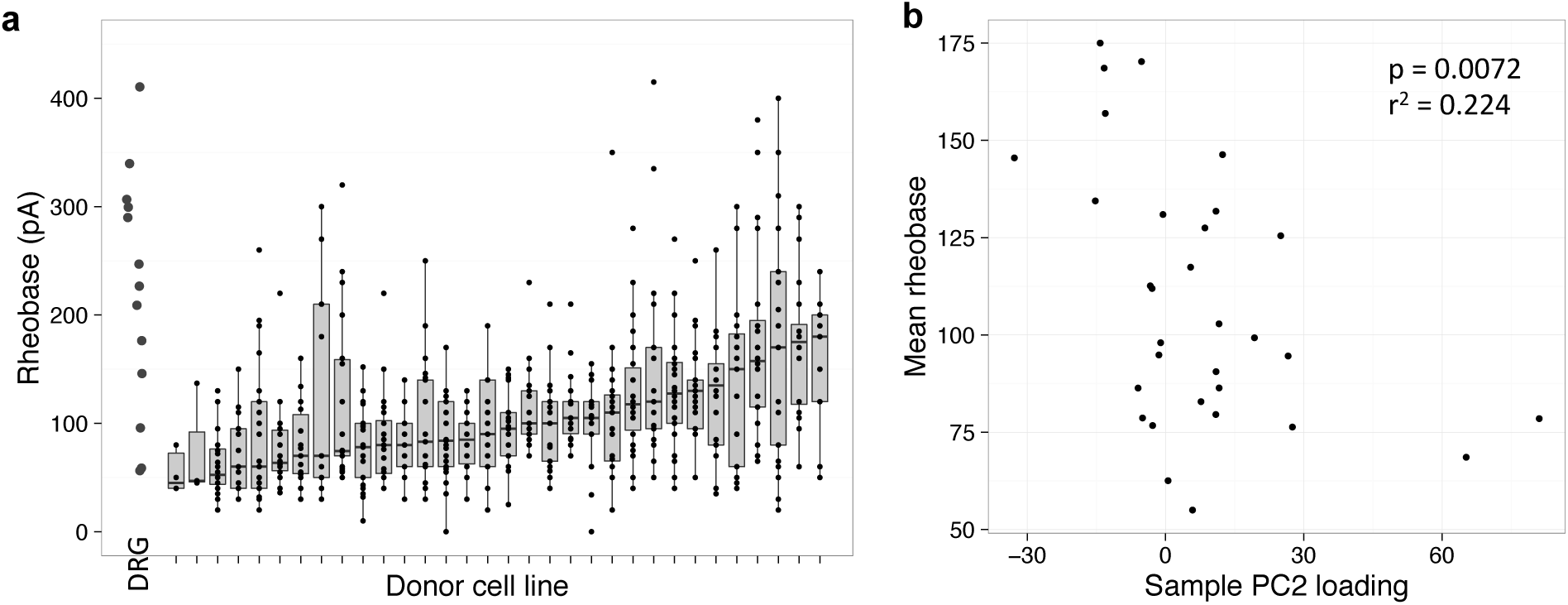
Functional profile and associations of IPSDSNs. (**a**) The distribution of rheobase values for the 31 samples with electrophysiology recordings. (**b**) Linear regression of principal component 2 of gene expression across samples with rheobase.

We next investigated whether variation in excitability was reflected in differences in gene expression of cells derived from the same donor. We examined the correlation between expression of individual genes and mean rheobase measured in sister cultures from the same donor and differentiation batch. After correcting for multiple testing, no individual genes were significantly correlated with rheobase at FDR < 0.1. However, we found a weak but significant correlation between the second principal component of gene expression, which accounted for 20.4% of expression variation, and rheobase (p=0.007, Pearson r^2^=0.22; Figure 3b), suggesting that there is a gene expression signature associated with variation in sensory neuronal excitability. That such an association was seen at all is striking, considering that the measurements were from separate flasks differentiated from IPSCs of a given donor, and one is a collection of single-cell measurements done after 6 weeks of differentiation while the other is a bulk sequencing assay done after 4 weeks of differentiation. Gene ontology analysis of genes with the top 500 positive and 500 negative PC2 loadings showed that samples with higher excitability (lower mean rheobase) were enriched for processes including neurogenesis and nervous system development, whereas samples with lower excitability were enriched for genes involved in extracellular matrix organization. For example, among the top three PC2 genes, *TTYH1* and *PTPRZ1* have known functions in nervous system development (Kleinman et al. 2014; Halleran et al. 2015; Kuboyama et al. 2015), while *NMU* encodes a multifunctional neuropeptide with a role in nociception (Martinez and O’Driscoll 2015). Among the bottom three PC2 genes (most negative loadings), *COL1A1* and *COL6A3* encode connective tissue collagen proteins, while *SERPINE1* is an inhibitor of fibrinolysis, and all three genes are more highly expressed in GTEx fibroblasts than any other GTEx tissue. Note that although PC1 accounts for 34% of the variation in gene expression and correlates with the fibroblast content of the samples, it did not correlate with rheobase (Supplementary Figure 17, p = 0.12). This is expected because only neuronal cells were measured by patch clamp, but it confirms that the presence of fibroblastlike cells did not significantly alter excitability of the sensory neurons.

### Genetic variants influence gene expression, splicing and chromatin accessibility in sensory neurons

We wished to discover expression quantitative trait loci (eQTLs) in sensory neurons, as these are candidate loci that may influence sensory neuron excitability and related complex traits such as pain sensation. However, variability in gene expression that is induced by the differentiation process could obscure genetic effects that are present. To address this, we used a recently developed method, RASQUAL (Kumasaka, Knights, and Gaffney 2016), which models both allele-specific and total expression to improve power for detecting cis-eQTLs. Allele-specific effects due to cis-acting genetic variants are robust to changes in a gene’s total expression due to *trans*-acting or non-genetic factors. After merging replicates and removing samples where we lacked genotype data, we had expression data for 97 unique donors. We tested the association of expression with all SNPs and indels with minor allele frequency >= 5% and within 500 kb of the transcription start site for 19796 protein-coding genes and 15237 noncoding RNAs. We adjusted for multiple testing at the gene level using Bonferroni correction based on the estimated number of independent tests for each gene as determined by EigenMT (Davis et al. 2016), and across genes by comparing with a single permutation of sample labels (see Methods).

At a genome-wide false discovery rate of 10% RASQUAL identified 3,778 genes with expression-modifying genetic variants, termed eGenes (Supplementary Table 3). In contrast, when using the linear model FastQTL with 10,000 permutations we identified only 1403 eGenes. The improvement in power was more striking when considering only significantly expressed genes (FPKM > 1), with RASQUAL and FastQTL discovering 2607 and 774 eGenes, respectively. Power gains relative to a linear model were greatest among genes with high variability across samples, and this was true for both novel associations and those reported previously in GTEx (Supplementary Figures 18,19). This illustrates the value of using an allele-specific model in the context of iPSC-derived cell types, where differentiation can lead to a mixture of cell types, causing increased gene expression variation as well as diluting signals of cell type-specific eQTLs.

Also of interest is whether we identify associations not already reported in GTEx (v6), the largest sequencing-based eQTL mapping study to date. For this we used a protocol described previously for the HIPSCI project (Kilpinen et al. 2016): we stringently designated an eQTL as tissue-specific when our lead SNP for a gene (or any of its high-LD proxies) did not have p < 2.2×10^−4^ for the same gene in any GTEx tissue (representing Bonferroni-corrected p <0.01 for 44 tissues). Notably, we did not observe significant eQTL sharing between IPSDSNs and the IPSCs they were derived from (Supplementary Figure 20a). Of all 3,778 eGenes, 954 had tissue-specific associations (Supplementary Table 6), including genes with known involvement in pain or neuropathies, such as *SCN9A, GRIN3A, P2RX7, CACNA1H/Cav3.2,* and *NTRK2.* The tetrodotoxin-sensitive fast voltage gated ion channel *SCN9A* in particular has been linked to both extreme pain disorders due to gain of function mutations in primary erythromelalgia (Cao et al. 2016), and congenital insensitivity to pain due to loss of function mutations (Cox et al. 2006). Because these eQTLs were not seen in any GTEx tissue, this suggests that these are regulatory variants with IPSDSN-specific function. In total we find eQTLs for 84 genes out of a set of 617 genes associated with pain from the literature (see Methods), of which 11 are IPSDSN-specific.

Variants affecting gene splicing (sQTLs) often change either protein structure or contextdependent gene regulation, and may be more enriched for complex trait loci than are eQTLs (Li et al. 2016). To detect sQTLs we used the annotation-free method LeafCutter (Li, Knowles, and Pritchard 2016) to define 59,736 clusters of alternatively spliced introns. After filtering to remove lowly-used intron junctions and clusters, we retained 30,591 clusters with on average 3.1 intron junctions per cluster. We then ran FastQTL (Ongen et al. 2016) with 10,000 permutations to associate intron usage levels with SNPs and indels within 15 kb of each intron’s midpoint. This yielded QTLs for 2,079 alternative splicing clusters at FDR 10% (Supplementary Table 4; see Methods). Notably, only 538 (26%) of the lead variants for these splicing associations were in linkage disequilibrium (LD) r^2^ >= 0.5 with a lead eQTL variant in our dataset, indicating that the sQTLs we identified further extend our catalog of expression-altering variants and are not merely proxies for gene-level eQTLs (or vice versa). In this regard, we found sQTLs for several pain-associated genes including *TRPV1, SCN3A, TAC1* and *P2RX4,* of which only *P2RX4* has an eQTL in our dataset.

Causal variants affecting gene expression and complex traits are highly enriched in regions of accessible chromatin (Degner et al. 2012), which can now be readily measured using ATAC-seq (Buenrostro et al. 2013). Moreover, these variants are often thought to act by altering binding of transcription factors to DNA, in turn affecting chromatin accessibility, looping of DNA to a gene promoter, and ultimately gene expression. Thus, causal variants altering chromatin accessibility (caQTLs) are expected to be found within the peaks themselves, which are typically small (< 1 kb). With ATAC-seq data for 31 samples and considering a small window of 2 kb centered on each peak (~380,000 peaks), we detected 6,318 caQTLs at FDR 10% in IPSDSNs (median of 6 variants tested per peak).

Although caQTLs do not themselves directly implicate a target gene, caQTLs overlapping eQTLs provide a hypothesis as to the causal regulatory variant altering a specific gene’s expression. We therefore considered all overlaps within 500 kb between lead variants for caQTLs and eQTLs. Surprisingly, of our 3778 eQTLs, only 410 were in LD (r^2^ >= 0.5) with a caQTL, with about half that number being in high LD (194 with r^2^ >= 0.8). We expected a larger overlap, as a recent estimate suggested that 50% of histone acetylation QTLs for H3K27ac affect expression of at least one gene within 500 kb (Li et al. 2016). Our lower overlap is likely due primarily to our requirement for both eQTL and caQTL to be independently genome-wide significant, as well as to our limited ATAC-seq sample size (31 samples). An alternative approach would be to test each caQTL for association with nearby genes, with the tradeoff that this would be much less stringent.

We sought to identify transcription factors in IPSDSNs whose binding is altered by the regulatory variants we identified. We used the LOLA Bioconductor package (Sheffield and Bock 2015) to test for enrichment of our lead QTL SNPs, relative to GTEx lead SNPs, in ENCODE ChIP-seq peaks and JASPAR transcription factor motifs. Considering first only our tissue-specific eQTLs, we found high enrichment within SMARCB1 and SMARCC2 peaks (odds ratios 5.8 and 14.1, respectively; p < 5×10^−5^), which are both members of the neuron-specific chromatin remodeling (nBAF) complex (Lessard et al. 2007). Also enriched were REST/NRSF (OR=5.7, p=1.1×10^-4^) and SIN3A (OR=3.9, p=1.0×10^−4^), which bind neuron-restrictive silencer elements during development, but have suggested roles in the development and maintenance of neuropathic pain (Willis et al. 2016). Extending the enrichment test to all IPSDSN eQTLs, we also found enrichments for ELK1 (OR=7.5, p=1.4×10^-13^) and ELK4 (OR=5.8, p=6.1x10^−16^), which are transcriptional activators with alternative splice isoforms expressed in neurons, and which co-bind DNA along with SRF (Kerr et al. 2010; Vanhoutte et al. 2001). Also enriched is c-Fos (OR=11.3, p=4.1×10^-14^), a target of ELK1 and ELK4, which is widely expressed but is known to have specific functions in sensory neurons (Hunt, Pini, and Evan 1987; Kohno et al. 2003). Notably, DNA sequence motifs for REST, ELK1 and ELK4 are also among the most highly enriched motifs in our ATAC-seq peaks. All enrichments are available in Supplementary Tables 7-9.

### Sensory neuron eQTLs and sQTLs overlap with complex trait loci

While we were interested in comparing our set of QTLs with GWAS for pain, the largest GWAS for pain to date included just 1,308 samples and found no associations at genome-wide significance (Peters et al. 2013). We therefore considered all GWAS catalog associations with p < 5×10^−8^ that were in high LD (r^2^ > 0.8) with a QTL in our dataset, with two purposes in mind: to determine whether any GWAS traits are enriched overall for overlap with sensory neuron QTLs, and to find individual cases where a QTL is a strong candidate as a causal association for the GWAS trait. IPSDSN eQTLs were significantly enriched for overlap with GWAS catalog SNPs (p < 0.001) relative to 1000 random sets of SNPs matched for minor allele frequency (MAF), distance to nearest gene, gene density, and LD (Pers, Timshel, and Hirschhorn 2014), and the overlap was consistent with that seen for eQTL studies in other tissues (Supplementary Figure 21). Although nociceptive neurons are specialized for sensing and relaying pain signals, they share characteristics with other neurons; thus, we might expect enrichment for traits known to involve the nervous system more generally. We restricted our analysis to the 41 traits with at least 40 GWAS catalog associations and then considered the binomial probability of overlap, with the expected overlap frequency being the proportion of QTL overlaps among all trait associations (6.2%). No traits showed significantly greater overlap with our QTL catalog than other traits after correcting for multiple testing (Supplementary Table 10).

Across all traits, we found 156 genes with an eQTL overlapping at least one GWAS association (Table 1), and similarly 129 sQTLs and 172 caQTLs with GWAS overlap; the full catalog of overlaps is reported in Supplementary Tables 11-13. We examined these associations, in conjunction with ATAC-seq peaks and LD information, to identify candidate causal variants influencing both a molecular phenotype and a complex trait.

**Table 1.** QTL associations. Columns show the number of associations, the number for genes associated with pain in the OpenTargets database, and the number of unique overlaps (r^2^ > 0.8) between lead QTL SNPs and GWAS catalog SNPs after removing duplicates for each GWAS trait.

|  | Number | Pain associated | GWAS overlap |
| --- | --- | --- | --- |
| eQTLs | 3778 | 84 | 156 |
| sQTLs | 2079 | 45 | 129 |
| ATAC QTLs | 6318 | N/A | 172 |
| Joint ATAC/eQTLs | 177 | 1 | 14 |

Among overlapping associations we find a number that relate to neuronal diseases, such as Parkinson’s disease, multiple sclerosis, and Alzheimer’s disease. One striking overlap is between an eQTL for SNCA, encoding alpha synuclein, and Parkinson’s disease, for which a likely causal variant has recently been identified (Soldner et al. 2016). The lead GWAS SNP and our lead eQTL are both in perfect LD with rs356168 (1000 genomes MAF 0.39), which lies in an ATACseq peak in an intron of SNCA. Soldner et al. used CRISPR/Cas9 genome editing in iPSCderived neurons to show that rs356168 alters both SNCA expression and binding of brain-specific transcription factors (Soldner et al. 2016). In IPSDSN cells we find that the G allele of rs356168 increases SNCA expression 1.14-fold, in line with Soldner et al. who reported 1.06- to 1.18-fold increases in neurons and neural precursors. However, despite residing in a visible ATAC-seq peak in our data, rs356168 is not detected as a caQTL (SNP p value = 0.22). eQTLs for SNCA have recently been reported in the latest GTEx release (v6p), but none of the tissue lead SNPs are in LD (r^2^ > 0.2) with rs356168, suggesting that the effect of this SNP can be more readily detected in specific cell and tissue types, including IPSDSNs and the frontal cortex tissue and iPSC derived neurons studied by Soldner et al.

A pain related trait of interest in this study is migraine. We find two QTLs overlapping GWAS associations from a recently reported meta-analysis of 375,000 individuals with migraine (Gormley et al. 2015). An eQTL for *ADAMTSL4* (rs1260387) is in high LD (r^2^ = 0.9) with a chromatin accessibility QTL at rs6693567, 11 kb upstream of the *ADAMTSL4* promoter. A number of lines of evidence suggest that of the two rs6693567 (MAF 0.30) is likely to be the causal variant in this region, although the causal gene is less certain. First, rs6693567 is the lead SNP for the migraine association, and is the only one among 8 SNPs in LD (r^2^ > 0.6, 1000 genomes phase 3 Europeans) which has substantial biochemical activity reported in ENCODE, where it sits in a peak of DNAse hypersensitivity in multiple cell types. It alters a conserved nucleotide (GERP score 4.75) in a TEAD4 ChIP-seq peak in H1-ES cells, 6 base pairs from a sequence motif for TEAD4. An eQTL at the same SNP is reported across many tissues in GTEx, and affects a number of genes in the region, including *ECM1, RP11-54A4.2, MRPS21, HORMAD1, GOLPH3L,* and *TARS2,* and in many cases rs6693567 is the lead SNP for those associations. It is not clear which of these genes is most relevant, although ECM1 is reported to have roles in angiogenesis (Han et al. 2001) and has been associated with psoriasis (Niu et al. 2016) and inflammatory bowel disease (Fisher et al. 2008). The most highly expressed of these genes in IPSDSNs is the mitochondrial ribosomal subunit MRPS21 consistent with reports of a link between mitochondrial dysfunction and migraine (Yorns and Hardison 2013).

We also find multiple compelling overlaps between splice QTLs and GWAS associations (Figure 4). One known example is a strong sQTL for *TNFRSF1A* (p=9.9×10^-29^) with the same lead SNP (rs1800693, MAF 0.30) as a multiple sclerosis association. This likely causal SNP is located 10 base pairs from the donor splice site downstream of exon 6, and has been experimentally shown to cause skipping of exon 6, resulting in a frameshift and a premature stop codon (Gregory et al. 2012). While *TNFRSF1A* is lowly expressed in neurons of the brain, it is highly expressed in many other tissues profiled by GTEx, including tibial nerve. We do not see an effect of this variant on total expression levels in our cells (p > 0.5), but we observe skipping of exon 6 in about 12% of transcripts from individuals homozygous for rs1800693 (Figure 4a). Since these transcripts undergo nonsense-mediated decay (Gregory et al. 2012), the actual rate of exon skipping is likely to be higher.

**Figure 4.**
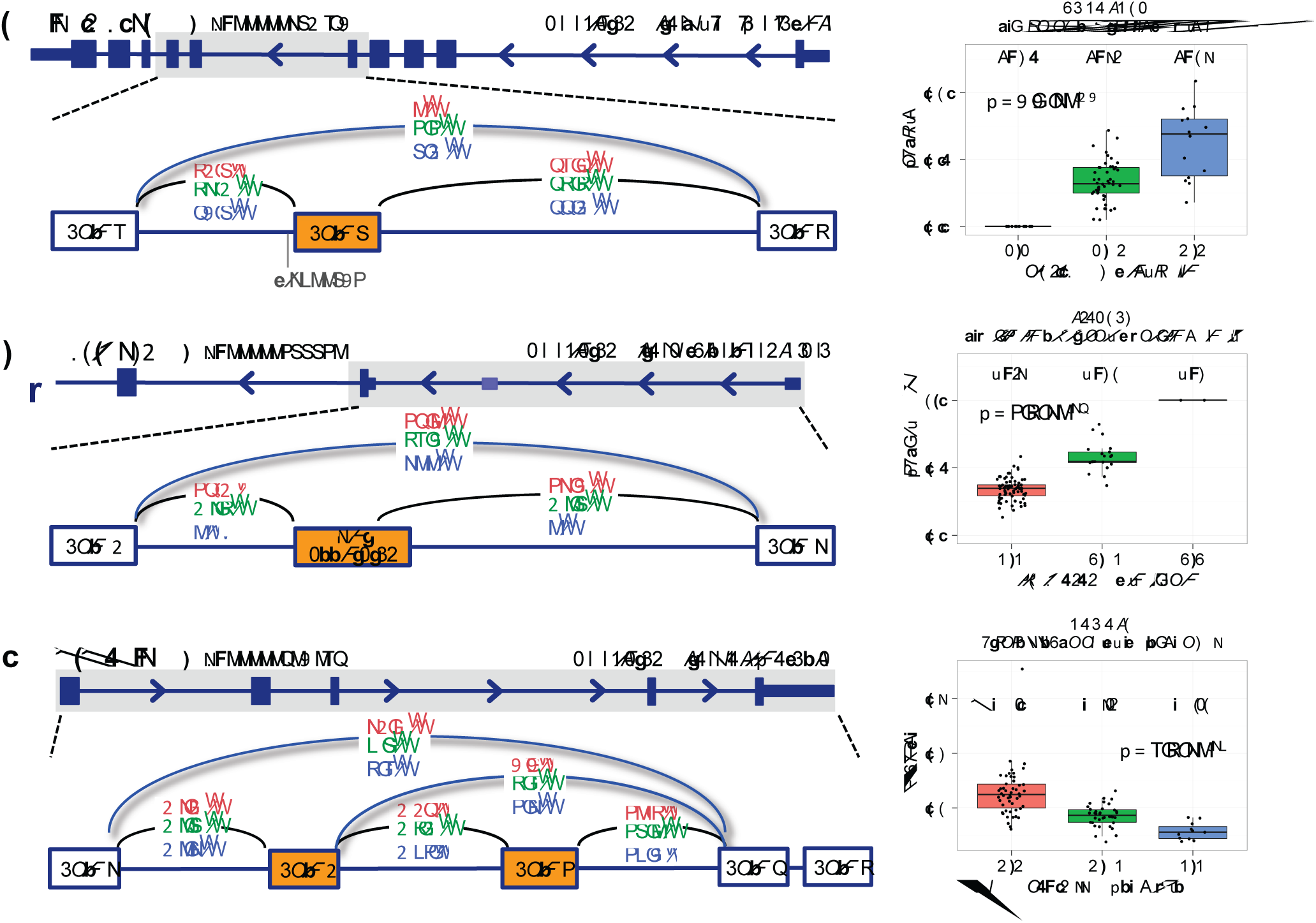
Splicing QTLs overlapping GWAS. **(a)** An sQTL for *TNFRSF1A* leads to skipping of exon 6, and overlaps with a multiple sclerosis association, **(b)** An sQTL for *SIPA1L2* leads to increased skipping of an unannotated exon between alternative promoters, and overlaps with a Parkinson’s disease association, **(c)** An sQTL for *APOPT1* alters skipping of exons 2 and 3, and overlaps with a schizophrenia association. P values are from the beta approximation based on 10,000 permutations as reported by FastQTL.

An sQTL for *SIPA1L2* (rs16857578, MAF 0.23) is in LD with associations for both Parkinson’s disease (rs10797576, r^2^=0.93) and blood pressure (rs11589828, r^2^=0.94). An unannotated noncoding exon (chr1:232533490-232533583) between alternative *SIPA1L2* promoters is included in nearly 50% of transcripts in individuals with the reference genotype, but splicing in of the exon is abolished by the variant (Figure 4b). SIPA1L2, also known as SPAR2, is a Rap GTPase-activating protein expressed in the brain and enriched at synaptic spines (Spilker and Kreutz 2010). Although its function is not yet clear, expression is seen in many tissues profiled by GTEx, with highest expression in the peripheral tibial nerve. Interestingly, the related protein SIPA1L1 exhibits an alternative protein isoform with an N-terminal extension that is regulated post-translationally to influence neurite outgrowth (Jordan et al. 2005).

A complex sQTL for *APOPT1* (rs4906337, MAF 0.22) is in near-perfect LD with a schizophrenia association (rs12887734). The splicing events involve skipping either of exon 3 only or both exons 2 and 3 (Figure 4c). At least 20 variants are in high LD (r^2^ > 0.9), including rs4906337 which is 40 bp from the exon 3 acceptor splice site, and rs2403197 which is 63 bp from the exon 4 donor splice site. No sQTL is reported in GTEx, and although eQTLs are reported for *APOPT1,* only the thyroid-specific eQTL (rs35496194) is in LD (r^2^ = 0.94) with the schizophrenia-associated SNP rs12887734. APOPT1 is localized to mitochondria and is broadly expressed. Homozygous loss-of-function mutations in this gene lead to Cytochrome c oxidase deficiency and a distinctive brain MRI pattern showing cavitating leukodystrophy in the posterior region of the cerebral hemispheres, with affected individuals having variable motor and cognitive impairments and peripheral neuropathy (Melchionda et al. 2014).

## Discussion

iPSC-derived cells enable the molecular mechanisms of disease to be studied in relevant human cell types, including those which are inaccessible as primary tissue samples. Because the effect sizes of common disease-associated risk alleles tend to be small, observing their effects in cellular models is challenging (Raghavan et al. 2016; Soldner et al. 2016). In an iPSC-based system, this difficulty is compounded by variability between samples in the success of differentiation, as described for hepatocytes (Dianat et al. 2013), hematopoietic progenitors (Smith et al. 2013), and neurons (Handel et al. 2016; Hu et al. 2010). Thus, for iPSC models of common disease associated variants to be meaningful, it is critical to know which candidate disease associated variants exhibit a cellular phenotype in an *in vitro* model.

Our study is the first that we are aware of to attempt iPSC differentiation at scale, from healthy individuals, and to functionally characterize the resulting differentiated cells and map the effects of common genetic variants genome-wide in an iPSC-derived cell type. We observed that, broadly, gene expression in our differentiated cells matched that observed in the closest primary tissue type we could obtain, dorsal root ganglia. However, we also observed that sample-to-sample variability in gene expression in the iPSC-derived cells was greater than that observed in DRGs. These highly variable genes were enriched in processes relating to neuronal differentiation and development. While this is unsurprising, it highlights that genes likely to be of particular interest and relevance for the function of these cells are also among the most variable, a challenge which may be broadly true of iPSC-derived cells. A portion of this variability is likely to be due to the imperfect process of *in vitro* differentiation, which produces a mixture of cell types.

Our single-cell RNA-seq data, which came from a single differentiation, could be cleanly clustered into neuronal cells and cells with more fibroblast-like gene expression. Using reference profiles from the two single-cell clusters enabled us to estimate the fraction of neuronal cells in our bulk RNA-seq samples, and these estimates qualitatively agreed with the neuronal content in images from the cell cultures. However, we do not have an independent quantitative measure of neuronal fraction, and we are mindful that no fixed clustering can fully represent the heterogeneity present in a cell population, whether iPSC-derived or from primary tissue. Our estimates of neuronal content in our bulk RNA-seq are thus dependent on the clusters we defined. The similarity that the fibroblast-like single cells also had with DRG raises the important question of whether these cells are immature sensory neurons, or are in a “confused” state that may not reflect any actual cells in the body. This question, which is relevant to all IPSC-derived cell types, could be addressed by single-cell sequencing done at multiple time points during differentiation. Alternatively, in neurons one can perform deep electrophysiological phenotyping followed by sequencing in the same single cells, known as Patch-seq (Cadwell et al. 2015) to enable classifying single neurons into functional categories and/or maturity levels. By establishing multiple reference profiles, these approaches could enable in silico sorting of cells based on their transcriptome, and better characterization of the sources of variation within a differentiated population of cells.

Neuronal excitability in IPSDSNs was correlated with the second principal component of gene expression, measured in separate cell cultures for a given donor and differentiation batch. Although this was a modest association (r2=0.224), it indicates that measurable differences in gene expression are recapitulated in higher-level neuronal phenotypes. Because neurons could be visually distinguished in our cultures, we were able to perform single-cell electrophysiology measurements only on apparently mature cells. Distinguishing mature differentiated cells from contaminating or immature cells might be more difficult in other derived cell types, in which case subsequent steps to isolate mature cells may be necessary (Choudhary et al. 2016). We did not detect a correlation between individual genes and neuronal excitability, likely due in part to the heterogeneity between individual neurons observed in each culture. When Patch-seq was recently used match transcriptional profiles and electrophysiology recordings in the same single cells, correlation between neuronal functional state and individual genes could be detected (Bardy et al. 2016).

Despite the observed sample-to-sample variability in gene expression, we detected thousands of eQTLs and caQTLs in IPSDSNs, boosting our power by incorporation of allele-specific information. Many of these overlap known expression-modifying variants that are associated with disease, including the eQTL for *SNCA* and Parkinson’s disease. Notably, many associations, including for *SNCA,* were discovered only with a model that statistically combines both allelespecific and between individual differences in expression to improve power for association mapping. Our results suggest that this approach is particularly useful in cases where there is significant sample-to-sample variability. We also discovered numerous QTLs for RNA splicing, using a method based on reads mapping across splice junctions (Li et al. 2016). Our catalog of QTLs includes a large number that overlap with complex trait associations, and for most of these the causal variants are not known. This QTL map is thus a starting point for in-depth dissection of individual loci in iPSC-derived cells where we have shown that a genetic effect is present.

In summary, we have measured multiple molecular phenotypes in the largest panel of iPSCderived cells to date. The catalog of QTLs we provide reveals a large set of common variants with detectable effects in IPSDSNs, suggesting promising targets for functional studies to fine-map causal disease-associated alleles in this cell type, such as by allelic replacement using CRISPRCas9.

## Acknowledgments

The iPSC lines were generated under the Human Induced Pluripotent Stem Cell Initiative (HIPSCI) funded by a grant from the Wellcome Trust and Medical Research Council, supported by the Wellcome Trust (WT098051) and the NIHR/Wellcome Trust Clinical Research Facility. HIPSCI funding was used for sensory neuron RNA-sequencing. We acknowledge Life Science Technologies Corporation as the provider of Cytotune. Pfizer Neuroscience (Pfizer Ltd.) funded neuronal differentiation, functional assays, single-cell RNA-sequencing, and collection and sequencing of dorsal root ganglion samples. The authors gratefully acknowledge Natsuhiko Kumasaka for help with RASQUAL. We thank Florian Merkle for comments on the manuscript. JS gratefully acknowledges support from the Wellcome Trust for his PhD studentship.

## URLs

OpenTargets, www.targetvalidation.org.

Cibersort, cibersort.stanford.edu.

ENCODE, www.encodeproject.org.

GTEx, www.gtexportal.org.

HIPSCI, www.hipsci.org.

## Data availability

RNA-seq and ATAC-seq for open access samples are deposited in the European Nucleotide Archive under accession XYZ. These data for managed access samples are deposited in the European Genome Archive under accession ZZZ. Sample genotypes and accession numbers are available at http://www.hipsci.org/data.

## Author contributions

JS analyzed data and wrote the manuscript. SF performed all differentiations. AGu analyzed data; AGu, DJG, and PJW conceived and supervised the project and revised the manuscript. HK compared eQTLs with GTEx and identified tissue-specific eQTLs. JR and MP cultured iPSC samples. AJK performed all ATAC-seq. KA and AGon assisted with data analysis. AW performed single cell RNA work and assisted with data analysis. RF and CLB performed RNA extraction and quantification. EI performed cell culture and Ca^2+^ flux assays. MB assisted with experimental design and Ca^2+^ flux assays. LC, SL, and AJL performed electrophysiology measurements. All authors reviewed the manuscript.

## Online methods

### IPS cell lines and neuron differentiation

Human induced pluripotent stem cells (iPSCs) from 103 healthy donors were obtained from the HIPSCI resource (Kilpinen et al. 2016). All samples for HIPSCI were collected from consented research volunteers recruited from the NIHR Cambridge BioResource (http://www.cambridgebioresource.org.uk). Samples were collected initially under existing Cambridge BioResource ethics for iPSC derivation (REC Ref: 09/H0304/77, V2 04/01/2013), with later samples collected under a revised consent (REC Ref: 09/H0304/77, V3 15/03/2013). All cell lines were reprogrammed to pluripotency using an identical protocol, as described for the HIPSCI resource. Of the 103 lines, 38 were initially grown in feeder-dependent medium and the remainder were grown in feeder-free E8 medium. Clump passaged iPSCs were single cell seeded in mTeSR1 (StemCell Technologies, Vancouver) media on hES-qualified Matrigel (BD Biosciences, San Jose, CA) 48 hours prior to neural induction (day 0). KSR media containing small molecule inhibitors LDN193189 (1μmol/l) and SB-431542 (10μmol/l) were added to cells from day 0 to 4 to drive anterior neuroectoderm specification. KSR Media was prepared as 500ml DMEM-KO (Life Technologies 10829-018), 130ml KSR-XF (Life Technologies 12618-013), 1x NEAA (Life Technologies 11140-068), 1x Glutamax (Life Technologies 35050-087), 0.01mM β-mercaptoethanol (Sigma M6250-100ml). From day 3, CHIR99021 (3μmol/l), DAPT (10μmol/l) and SU5402 (10 μmol/l) were added to further enable the emergence of neural crest phenotypes. N2B27 media was used incrementally every two days from D3. N2B27 Media was prepared as 500ml Neurobasal medium (Life Technologies 21103-049), 5ml N2 supplement (Life Technologies 17502-048), 10ml B27 supplement without vitamin A (Life Technologies 12587-010), 0.01mM β-mercaptoethanol (Sigma M6250-100ml) and 1x Glutamax (Life Technologies 35050-087). On day 11 maturation media containing N2B27 media with human-b-NGF (25ng/ml), BDNF (25 ng/ml), NT3 (25ng/ml) and GDNF (25ng/ml) was and used for long term culture. AraC treatment 4μM) was used once at day 14 to reduce the non-neuronal population when necessary. Cells were differentiated in T25 flasks for RNA and nuclei isolation and replated at day 14 onto coverslips and 96 well plates for electrophysiology and Ca^2+^ flux assays.

### Single-cell RNA sequencing

Single cells were sequenced from a single sample. Although this sample used the same differentiation protocol, it was not derived from a HIPSCI donor and it was matured for 8 weeks, whereas the RNA-seq samples were matured 4 weeks. Previous work showed only minor changes in gene expression between 4 and 8 weeks maturation (Young et al. 2014). Dissociated cells were loaded onto a Fluidigm C1 system for automatic cell separation, reverse transcription and amplification. Libraries were only prepared from C1 chambers that contained single cells, using the Illumina Nextera XT kit as per the Fluidigm C1 protocol. These were quantified with the Qubit dsDNA HS assay (Thermo Fisher) and KAPA Library Quantification Kit (KAPA Biosystems) and size-checked with the Agilent Bioanalyser DNA 1000 assay (Agilent), as per manufacturers’ recommendations. Libraries were 96-way multiplexed and sequenced paired end on an Illumina Nextseq500 (75bp reads). Reads for each cell were aligned to GRCh38 and Ensembl 80 transcript annotations using STAR v2.4.0d with default parameters.

We had gene expression counts for ~56,000 genes (including noncoding RNAs) for 186 cells, although many of these were zeros. We excluded 9 cells expressing fewer than 20% of the quantified genes, and then used SC3 (Kiselev et al. 2016) to cluster the remaining 177 cells based on expression counts. Note that when clustering cells from complex tissues there is often a hierarchy of clusters, and no specific number of clusters can be considered correct. Allowing that the same could be true of IPS-derived cells, we examined alternative numbers of clusters from k=2 to 5 (Supplementary Figure 6), specifying k (the number of clusters) ranging from 2 to 5. With two clusters, the marker genes reported by SC3 clearly identified one cluster (111 cells) as neuronal, whereas the other cluster (66 cells) had high expression of extracellular matrix genes reminiscent of fibroblasts.

To display marker gene expression we selected 5 neuronal and 5 fibroblast marker genes based on the literature. After DESeq2’s variance stabilizing transformation, we used R’s “scale” function to mean-center and normalize expression values across cells for these genes, and plotted the result using the pheatmap R package.

With 3 and 4 clusters, the sensory-neuronal cell cluster remained unchanged, and the fibroblastlike cluster became further subdivided. This suggests that a majority of the cells in this sample were terminally differentiated into sensory neurons, whereas the remaining cells were more heterogeneous in their gene expression. With 5 clusters, the neuronal cluster split in two, while the fibroblast-like clusters remained unchanged.

### Genotypes

We obtained imputed genotypes for all of the samples from the HIPSCI project. We used CrossMap (http://crossmap.sourceforge.net/) to convert variant coordinates from GRCh37 reference genome to GRCh38. We then used bcftools (http://samtools.github.io/bcftools/) to retain only bi-allelic variants (SNPs and indels) with INFO score > 0.8 and MAF > 0.05 in the 97 samples used for QTL calling. This filtered VCF file was used for all subsequent analyses.

### RNA sequencing

Cells growing in T25 flasks were washed twice with PBS followed by addition of 600 mL of RLTPlus buffer. Cells were gently lifted from the flask and transferred to 1.5 ml tubes. Lysates were transferred to 1.5 mL tubes. RNA and gDNA were isolated using AllPrep DNA/RNA Minikit (Qiagen). RNA was eluted in 33 uL of DNAse free water and DNA eluted in 53 uL EB buffer.

RNA libraries were prepared using the Illumina TruSeq strand-specific protocol, and were sequenced with paired-end reads (2x75) on Illumina HiSeq with V4 chemistry. There were 131 RNA samples, which corresponded with 103 unique HIPSCI cell lines, as some of the samples were differentiation replicates or RNA-extraction replicates. One sample failed in sequencing and was excluded. Reads for each sample were aligned to GRCh38 and Ensembl 79 transcript annotations using STAR v2.4.0j with default parameters. We used VerifyBamID v1.1.2 (Jun et al. 2012) to check that RNA-seq sample BAM files matched the corresponding sample genotypes in the core HIPSCI VCF files. This revealed 5 mislabeled RNA samples, for which the correctly matching sample genotypes could be easily determined and corrected, as well as two samples for which no match could be found in HIPSCI genotype data and which were thus excluded (these had been labeled as problematic samples in HIPSCI).

### ATAC library preparation and sequencing

#### Nuclei isolation

Media was removed from T25 flasks and washed twice with 10 mL of room temperature D-PBS without calcium and magnesium. The adherent neuronal cultures were lifted by treating with 3 mL of Accutase (Millipore – SCR005) at room temperature for four minutes. The Accutase was quenched by adding 6 mL of 2 % foetal bovine serum in D-PBS. The cells were transferred to a 15 mL conical tube and centrifuged at 300 g for 5 minutes at 4 °C. The cell pellet was resuspended in 1 mL of ice-cold sucrose buffer (10 mM tris-Cl pH 7.5, 3 mM CaCl_2_, 2 mM MgCl_2_ and 320 mM sucrose) and pipetted briefly to break up the large clumps before incubating on ice for 12 minutes. 50 μL of 10% Triton-X 100 was added to the sucrose-treated cells and mixed briefly before incubating on ice for a further 6 minutes. Nuclei were released by performing 30 strokes with a tight dounce homogeniser on ice. Approximately 1 × 10^5^ nuclei were transferred to a 1.5 mL microfuge tube and centrifuged at 300 g for 5 minutes at 4 °C. All traces of the lysis buffer were removed from the nuclei pellet.

#### Tagmentation, PCR amplification and size selection

The tagmentation and PCR methods used here are in principle the same as that described in Buenrostro et al., 2013, but with some modifications as described in Kumasaka et al., 2016. The nuclei pellet was resuspended in 50 μL of Nextera tagmentation master mix (Illumina FC-121-1030) (25μL 2x Tagment DNA buffer, 20 μL nuclease-free water and 5 μL Tagment DNA Enzyme 1) and incubated at 37 °C for 30 minutes. The tagmentation reaction was stopped by the addition of 500 μL Buffer PB (Qiagen) and purified using the MinElute PCR purification kit (Qiagen 28004), according to the manufacturer’s instructions and eluting in 10 μL of Buffer EB (Qiagen). 10 μL of the tagmented chromatin was mixed with 2.5 μL Nextera PCR primer cocktail and 7.5 μL Nextera PCR mastermix (Illumina FC-121-1030) in a 0.2 mL low-bind PCR tube. The indexing primers used for amplification were from the Nextera Index kit (Illumina F C-121-1011), using 2.5μL of an i5 primer and 2.5μL of an i7 primer per PCR, totalling 25μL. PCR amplification was performed as follows: 72 °C for 3 minutes and 98 °C for 30 seconds, followed by 12 cycles of 98°C for 10 seconds, 63 °C for 30 seconds and 72 °C for 3 minutes. To remove the excess of unincorporated primers, dNTPS and primer dimers, Agencourt AMPure XP magnetic beads (Beckman Coulter A63880) were used at a ratio of 1.2 AMPure beads:1 PCR sample (v/v), according the manufacturer’s instructions, eluting in 20μL of Buffer EB (Qiagen). Finally, size selection was performed by 1% agarose TAE gel electrophoresis, selecting library fragments from 120 bp to 1 kb. Gel slices were extracted with the MinElute Gel Extraction kit (Qiagen 28604), eluting in 20μL of Buffer EB.

#### Illumina sequencing

A total of 31 ATAC-seq libraries each prepared with a unique Nextera i5 and i7 tag combination were pooled. Index tag ratios were assessed by a single MiSeq run and were balanced before being sequenced at two per lane with paired-end reads (2×75) on a HiSeq with V4 chemistry. However, rebalancing did not appear to work correctly, as the number of reads varied greatly between samples, from a minimum of 17 million to a maximum of 987 million. However, 22 samples had over 100 million reads, and 30 samples had over 40 million reads. Across samples, a median of 56% of reads mapped to mitochondrial DNA. For calling ATAC QTLs we used all sample counts as-is.

#### Read alignment

We aligned reads to GRCh38 human reference genome using bwa mem v0.7.12. Reads mapping to the mitochondrial genome and alternative contigs were excluded from all downstream analysis. As for RNA-seq data, we used VerifyBamID v1.1.2 (Jun et al. 2012) to detect sample swaps. This revealed one mislabeled sample, which we then corrected. We used Picard v1.134 MarkDuplicates (https://broadinstitute.github.io/picard/) to mark duplicate fragments. We constructed fragment coverage BigWig files using bedtools v2.21.0 (Quinlan and Hall 2010).

#### Peak calling

We used MACS2 v2.1.1 (Zhang et al. 2008) to call ATAC-seq peaks individually on sample BAM files with parameters ‘--nomodel --shift -25 --extsize 50 -q 0.01’. We then constructed a consensus set of peaks by determining regions in which peaks overlapped in at least 3 samples. At regions of overlap, the consensus peak was defined as the union of overlapping peaks. This resulted in 381,323 peaks, with 98% of peaks ranging in size from 82 – 1191 base pairs.

### Gene expression quantification, quality control and exclusions

GTF files for the Gencode Basic transcript annotations, GRCh38 release 79, were downloaded from www.gencodegenes.org. We excluded short RNAs, pseudogenes, and genes not mapping to chromosomes 1–22, X, Y, or MT, leaving 35,033 unique genes. Gene expression counts were determined using the featureCounts tool of the subread package v1.5.0 (Liao, Smyth, and Shi 2014) with options (-s 2 -p -C -D 2000 -d 25). A median of 45 million reads were generated per sample, with median 32.8 million reads (72%) uniquely mapping and assigned to genes. Expression counts were normalized using conditional quantile normalization with the R package cqn v5.0.2 (Hansen, Irizarry, and Wu 2012). We defined expressed genes as the 14,215 genes with mean CQN-normalized expression across samples > 1.

We determined pairwise correlation between samples using normalized counts for expressed genes and plotted these as a heatmap. We also plotted the first five principal components of gene expression against each other. These plots identified four outlier samples, which were excluded from subsequent analyses (Supplementary Figure 1). After all exclusions and corrected sample labels, we retained 126 samples from 99 unique donors. For gene expression quantification and QTL calling (both eQTL and sQTL), replicate BAM files from same donor were merged together using samtools. Although this meant that some samples had about three times as many reads as other samples, neither of the QTL calling methods we used is strongly sensitive to this difference. Because genotypes were not available from HIPSCI for two donors, we retained gene expression data for 97 donors for QTL calling.

#### Assessing gene expression replicability

We used R with ggplot2 to plot the CQN-normalized expression for pairs of sample replicates.We excluded 16 samples differentiated using the first version protocol (P1), as most samples (110) were differentiated with the second version (P2), which gave us sufficient samples to consider variability between differentiations without including protocol effects. We determined the spearman correlation coefficient across all genes for (a) extraction replicates, (b) differentiation replicates, and (c) all possible pairs of samples from different donors. The histogram of correlation coefficients for these 3 categories is shown in Supplementary Figure 2.

### Dorsal root ganglion samples and sequencing

Human tissue acquisition and handling was performed at Pfizer Neuroscience in accordance with regulatory guidelines and ethical board approval. Postmortem human dorsal root ganglia (DRG) were obtained in dissected form from Anabios or as an encapsulated sheath together with sensory/afferent axons from National Disease Research Interchange which were subsequently dissected to isolate the cell-body rich ganglion. The tissue was homogenized in an appropriate volume QIAzol Lysis Reagent according to weight and processed according to the manufacturer's instructions for the Qiagen RNeasy Plus lipid-rich kit. RNAseq library preparation and sequencing was performed using the Illumina TruSeq Stranded mRNA Library Prep Kit and an Illumina HiSeq 2500 generating 2 x 100 bp reads by Aros Inc. according to the manufacturer’s instructions. Sequencing reads were aligned to the GRCh37 reference human genome using STAR and gene counts and FPKMs obtained using featureCounts and Ensembl v80 gene annotations.

### Multidimensional scaling plot clustering samples with GTEx tissues

We downloaded the GTEx v6 gene RPKM file (GTEx_Analysis_v6_RNA-seq_RNA-SeQCv1.1.8_gene_rpkm.gct.gz) as well as sample metadata (GTEx_Data_V6_Annotations_SampleAttributesDS.txt) from the GTEx web portal (http://www.gtexportal.org/home/datasets). We computed RPKMs for all genes for the 28 DRG samples, the 119 sensory neuron samples (5 outliers removed), and all HIPSCI IPS samples. We used all genes that were quantified in all of these sample sets. We determined pairwise sample distances in R (d = 1 - cor(rpkm.matrix)), then computed MDS locations (isoMDS(d, k=2)) with MASS package and plotted the results with ggplot2.

### Highly variable genes in IPSDSNs and GTEx

We obtained GTEx v6 RPKM files for all genes as described above. For each of the 44 tissues, as well as IPSDSNs, DRG, and HIPSCI IPSCs, we calculated the coefficient of variation (CV) of each gene among samples with the same detailed tissue type (SMTSD in GTEx sample metadata). We then subsetted the genes considered in each tissue to those expressed at RPKM > 1 in that tissue. We plotted the distribution of CVs across all genes for each tissue as a density plot (Supplementary Figure 3). We also calculated “CVRatio” for each gene in each tissue, which is the ratio of the gene’s CV to the mean CV for the gene in all other tissues where the gene is expressed. As an example, suppose that gene A has a CV of 0.6 in tissue X, and is expressed in three other tissues where it has CVs of 0.2, 0.3, and 0.7. The CVRatio for gene A in tissue X would be 0.6 / ((0.2 + 0.3 + 0.7) / 3) = 1.5. This is meant to indicate how variable gene A’s expression is in tissue X relative to its variability in other tissues, i.e. to identify “unusually variable” genes. In practice, the difference in gene rankings between CV and CVRatio for IPSDSNs was not large, although slightly more genes could be considered outliers in IPSDSNs with CVRatio.

We used the GOSummaries R package (Kolde and Vilo 2015) to compute gene ontology enrichments for the top 1000 most highly variable genes in IPSDSNs by CVRatio, which are reported in Supplementary Table 2. GOSummaries uses the g:Profiler web tool for enrichments (http://biit.cs.ut.ee/gprofiler/) (Reimand, Arak, and Vilo 2011). GO enrichments were similar whether considering the top 1000 genes by CV or by CVRatio.

### Estimation of neuronal purity

We used Cibersort (Newman et al. 2015) to estimate the fraction of RNA from neuronal cells in our bulk RNA-seq samples. We used the 14,786 genes whose CQN expression in bulk RNA samples was greater than zero, and retrieved raw counts for these genes in our single cell RNA-seq data. We labeled the single cells as “neuron” or “fibroblast-like” as determined based on the SC3 clustering, and specified these single cell counts as the reference samples for Cibersort to generate a custom signature genes file during its analysis. We used raw expression counts for the same genes for our 126 bulk RNA-seq samples as the mixture file for Cibersort to use in estimating the relative fractions of neuron and fibroblast-like cell RNA.

### Electrophysiological recordings

Six coverslips per line were placed singularly into a 12-well plate and washed 1x with 1 ml DPBS (+/+). After removal of DPBS, the coverslips were coated with 1 ml of 0.33 mg/ml growth factor reduced matrigel for > 3 hr at room temperature. D14 cells were prepared at a suspension of 1.6x10^6^/ml in 15 ml media. The cells were then diluted in NB media to create a 0.3e6/ml suspension. The coverslips were transferred into a clean 12-well plate and 1 ml of the cell suspension was added. Plates were incubated at 37°C (5% CO2) in a cell culture incubator for 24hrs, after which the coverslips were transferred into a clean 12-well plate containing 2 ml media. Cells were then treated with Mitomycin C (0.001 mg/ml for 2hr hours at 37°C) post plating on day 4 and day 10. Media was changed twice weekly.

Patch-clamp experiments were performed in whole-cell configuration using a patch-clamp amplifier 200B for voltage clamp and Multiclamp 700A or 700B for current clamp controlled by Pclamp 10 software (Molecular Devices). Experiments were performed at 35°C or 40°C as noted controlled by an in-line solution heating system (CL-100 from Warner Instruments). Temperature was calibrated at the outlet of the in-line heater daily before the experiments. Patch pipettes had resistances between 1.5 and 2 MΩ. Basic extracellular solution contained (mM) 135 NaCl, 4.7 KCl, 1 CaCl_2_, 1 MgCl_2_, 10 HEPES and 10 glucose; pH was adjusted to 7.4 with NaOH. The intracellular (pipette) solution for voltage clamp contained (mM) 100 CsF, 45 CsCl, 10 NaCl, 1 MgCl_2_, 10 HEPES, and 5 EGTA; pH was adjusted to 7.3 with CsOH. For current clamp the intracellular (pipette) solution contained (mM) 130 KCl, 1 MgCl_2_, 5 MgATP, 10 HEPES, and 5 EGTA; pH was adjusted to 7.3 with KOH. The osmolarity of solutions was maintained at 320 mOsm/L for extracellular solution and 300 mOsm/L for intracellular solutions. All chemicals were purchased from Sigma. Currents were sampled at 20 kHz and filtered at 5 kHz. Between 80% and 90% of the series resistance was compensated to reduce voltage errors. The voltage protocol used for the compounds testing on voltage gated sodium channels consisted of steps from a holding potential of -110 mV to -70 mV for 5 seconds, followed by step to -110 mV for 100 millisecond then currents were measured at step to 0 mV for 20 milliseconds. Intersweep intervals were 15 seconds. Rheobase was measured in current clamp mode by injecting increasing 30 milliseconds current steps until a single action potential was evoked. Intersweep intervals were 2 seconds. Membrane potential was set at either free-resting or held at -70 mV as noted. Current clamp data was analyzed using Spike2 software (Cambridge Electronic Device, UK) and Origin 9.1 software (Originlab).

### Correlation of gene expression with electrophysiological phenotypes

To avoid double-counting samples, we selected the first sample from each differentiation replicate to include in the gene expression table (104 samples). We then used DESeq2’s variance stabilizing transformation on the raw gene expression counts. We computed the first 10 principal components (PCs) with Bioconductor’s pcaMethods package. The first 3 PCs accounted for 34%, 20%, and 7% of expression variability, respectively, whereas subsequent PCs explained <= 3% of variability each. We therefore considered only the first 3 PCs in subsequent analyses. For the 31 samples with rheobase measurements, we examined the Pearson correlation between samples’ mean rheobase values and their PC loadings. Only PC2 showed a significant correlation upon linear regression (p=0.0072, Pearson r^2^=0.22).

We also examined correlation of individual genes with rheobase. We used DESeq2 with PC1 and mean rheobase in the design formula, and excluded genes with a maximum Cook’s outlier score above 5. At FDR 10%, DESeq2 reported 19 genes whose expression correlated with rheobase. However, on further examination these associations still appeared to be driven by outlier samples. We did a similar analysis considering spearman correlation between mean rheobase and each gene’s expression across samples, but no genes had a significant correlation at any FDR threshold when considering the number of tests done.

### QTL calling

#### Expression QTLs

To call cis-eQTLs we used RASQUAL (Kumasaka, Knights, and Gaffney 2016), which leverages allele-specific reads in heterozygous individuals to improve power for QTL discovery, while accounting for reference mapping bias and a number of other potential artifacts. With RASQUAL a feature is defined by a set of start and end coordinates; for calling a gene eQTL these are the start and end coordinates for exons, whereas for an ATAC-seq peak these are the peak coordinates. RASQUAL requires as input the allele-specific read counts at each SNP within a feature. We used the Genome Analysis Toolkit (GATK) program ASEReadCounter (Castel et al.2015). with options ‘-U ALLOW_N_CIGAR_READS -dt NONE --minMappingQuality 10 -rf MateSameStrand’ to count allele-specific reads at SNPs (and not indels). We then annotated the AS read counts in the INFO field of the VCF used as input for RASQUAL. We used custom scripts to determine the number of feature SNPs in gene exons.

We used RASQUAL’s makeCovariates.R script to determine principal components (PCs) to use as covariates, which determined 12 PCs as appropriate from the expression count data. We ran RASQUAL separately for each of 35,033 genes, passing in VCF lines for all SNPs and indels (MAF > 0.05, INFO > 0.8) within 500 kb of the gene transcription start site. To correct for multiple testing we used permutations; however, because RASQUAL is computationally intensive, it would not be possible to run a thousand or more permutations for every gene. Therefore we used an approach to balance power and computational time. To correct for the number of SNPs tested per gene, we used EigenMT (Davis et al. 2016) to estimate the number of independent tests per gene, and then performed Bonferroni correction on a gene-by-gene basis. To estimate the false discovery rate (FDR) across genes, we used the --random-permutation option of RASQUAL and re-ran it once for every gene, saving the minimum p value (after EigenMT correction) of the SNPs tested for each gene. This gave a distribution of minimum p values across genes for the permuted data. To determine the FDR for eQTL discovery at a given gene, we use R to compute (#permuted data min pvalues <p) / (#real data min p values < p), where p is the minimum p value among SNPs for the gene in question. With this procedure we obtained 3,586 genes with a cis-eQTL at FDR 10% (2,628 at FDR 5%).

#### ATAC QTLs

As we did for gene expression, we used featureCounts v1.5.0 to count fragments overlapping consensus ATAC-seq peaks and ASEReadCounter to count allele-specific reads at SNPs (and not indels) within peaks. We ran RASQUAL separately for each of 381,323 peaks, passing in VCF lines for SNPs and indels (MAF > 0.05, INFO > 0.8) within 1 kb of the center of the peak. Since >99.9% of peaks were less than 2 kb in size, this meant that we tested effectively all SNPs within peaks. As we did when calling eQTLs, we ran RASQUAL with the --random-permutation option for every gene, and determined FDR as described above. Note that in this case we used Bonferroni correction based on the number of SNPs tested, without using EigenMT, due to the small size of the windows tested. With this procedure we obtained 6,318 ATAC peaks with a cisQTL at FDR 10%.

#### Splice QTLs

We downloaded LeafCutter from Github (https://github.com/davidaknowles/leafcutter) on April 17, 2016. We used the LeafCutter bam2junc.sh script to determine junction counts for each sample, followed by leafcutter_cluster.py. This resulted in 254,057 junctions in 59,736 clusters. To focus on splicing events likely to be significant, we applied a number of filters, including: (a) removing junctions accounting for less than 2% of the cluster reads, (b) removing introns used (i.e. having at least 1 supporting read) in fewer than 5 samples, (c) retaining only clusters where at least 10 samples had 20 or more reads in the cluster. This yielded a filtered set of 95,786 junctions in 30,591 clusters. We first determined the read proportions for all junctions within alternatively excised clusters. We then Z-score standardized each junction read proportion across samples, and then quantile-normalized across introns. We used this as our phenotype matrix for input to FastQTL to test for associations between intron usage and variants within 15 kb of the center of each intron. We chose a cis-window size of 30 kb (2 × 15 kb) because >91% of introns are < 30 kb in size, and so this tests variants near exon/intron boundaries for the great majority of introns, while maximizing power.

We ran FastQTL in nominal pass mode 31 times specifying the first 0 to 30 principal components as covariates, and examined the number of intron QTLs with minimum SNP p value < 10^-5^. This showed that the number of QTLs plateaued when 5 PCs were used, and so we used 5 PCs in subsequent runs. We next ran FastQTL with 10,000 permutations to determine empirical p values for each alternatively excised intron. To correct for the number of introns tested per cluster, we used Bonferroni correction on the most significant intron p value per cluster. We then used the Benjamini-Hochberg method to estimate FDR across tested clusters. This yielded 2,079 significant SNP associations for intron usage (sQTLs) at FDR 10%.

For significant sQTLs we used bedtools closest with GRCh38 release 84 to annotate the gene(s) nearest the lead SNP for the association. To ensure we had relevant genes, we filtered the annotation to include only genes where one of the exon boundaries matched the intron boundary for the sQTL.

### Identifying tissue-specific eQTLs

We determined the set of tissue-specific eQTLs using the same procedure and code as in the HIPSCI project (Kilpinen et al. 2016). Briefly, we considered the full cis eQTL output of sensory neuron eQTLs and 44 tissues analyzed by the GTEx Project (Mele et al. 2015). For each discovery tissue (including sensory neurons), we tested for the replication of all lead eQTL -target eGene pairs reported at FDR 5%. If the lead eQTL variant was not reported in the comparison tissue, then the best high-LD proxy of the lead variant (r^2^ > 0.8 in the UK10k European reference panel) was used as the query variant. Replication was defined as the query variant having a nominal eQTL p < 2.2×10^-4^ (corresponding to p = 0.01 / 45, where 45 refers to the total number of tissues tested) for the same eGene. We then extracted eGenes for which the lead eQTL did not show evidence of replication in any other tissue (p > 2.2×10^−4^) or could not be tested (i.e. was not measured or reported as expressed in any other tissue).

This analysis gave 954 eGenes where the eQTL is specific to sensory neurons (Supplementary Table 6). We note that some of these “tissue-specific” eGenes could be due to the difference in QTL-calling methods used, notably that we used RASQUAL, a method incorporating both allelespecific and population-level expression variation. Therefore, some of the tissue-specific eGenes we report may actually be present more broadly in GTEx tissues but missed by the linear QTL model used in GTEx. Among the 1403 eGenes called by FastQTL, 208 were tissue-specific to IPSDSNs.

### Pain-associated genes

We identified a set of pain-associated genes by searching for the term “pain” in the OpenTargets web site (https://www.targetvalidation.org/) on August 22, 2016, and downloading the reported gene associations and scores. We chose a score cutoff of 0.05 to designate a gene as pain-associated, which resulted in 617 genes.

### Motif enrichment analyses

We used the R Bioconductor package LOLA (Sheffield and Bock 2015) to identify enrichments in transcription factor binding sites (TFBS) and motifs. We defined three sets of loci to consider for enrichment: 1) tissue-specific eQTL SNPs with a window of 50 bp (+/-25) around the SNP position, 2) all eQTL SNPs (50 bp window), and 3) all ATAC-seq peaks. For the QTLs we used all GTEx eQTL lead SNPs as the “universe” set against which we were testing TFBS for enrichment. For this we downloaded all GTEx QTL files (*_Analysis.snpgenes), loaded them in R and used the liftOver function from the rtracklayer package to convert their coordinates to the GRCh38 genome version. We tested for enrichment against the LOLA core database but considered only ENCODE TFBS enrichments. These enrichments are reported in Supplementary Tables 7 and 8. We also tested for enrichment against the LOLA extension database and considered JASPAR motif enrichments. No motif enrichments were found for IPSDSN eQTLs relative to GTEx eQTLs. We also tested ATAC-seq peaks for enrichment relative to DNase hypersensitive sites for many tissues from Sheffield et al. (Sheffield et al. 2013), which are available in the LOLA catalog. Many of the same TFBS enrichments were seen for ATAC-seq peaks as for eQTLs (data not shown), although with a skew towards general transcription factors (e.g. CTCF, ATF3, MYC, JUN) as might be expected. Motif enrichments in ATAC-seq peaks are reported in Supplementary Table 9.

### QTL overlap with GWAS catalog

The GWAS catalog was downloaded from https://www.ebi.ac.uk/gwas/ on 2016-5-08. To determine overlap between variants in the GWAS catalog and our lead QTLs, we first extracted all lead variants (both QTLs and GWAS catalog variants) from the full VCF file. We used vcftools v0.1.14 (Danecek et al. 2011) to compute the correlation R^2^ between all lead variants within 500 kb of each other among our samples. We determined overlap separately for eQTLs, sQTLs, and ATAC QTLs, and retained only overlaps with R^2^ > 0.8 between lead variants. Note that a given GWAS variant may be in LD with an eQTL for more than one gene, and vice versa, an eQTL for a single gene may be in LD with more than one GWAS catalog entry.

We used QTL-GWAS overlap for two purposes: first, to find individual cases where a QTL is a strong candidate as a causal association for the GWAS trait, and second, to determine whether any GWAS catalog traits are enriched overall for overlap with sensory neuron QTLs. For the first goal, we considered all overlaps with GWAS catalog associations having p < 5×10^-8^, i.e. did not filter any redundant overlaps. These overlaps are reported in Supplementary Tables 11 (for eQTLs), 12 (for sQTLs), and 13 (for ATAC QTLs).

To determine whether our QTL overlaps were enriched in any specific GWAS catalog traits relative to other traits, we computed overlap with all GWAS catalog SNPs (p < 5×10^−8^) but sought to eliminate redundant overlaps. For traits that were reported with differing names (e.g. “Alzheimer’s disease (cognitive decline)” and “Alzheimer's disease in APOE e4-carriers”), we grouped these into a single trait name (e.g. “Alzheimer’s disease”). We then sorted overlaps by decreasing LD R^2^, and kept the single overlapping QTL with the highest R^2^ for each GWAS catalog entry. Similarly, we removed duplicates with the same reported GWAS catalog SNP and trait, such as when successive GWAS of the same trait report the same SNP association. We counted the number of such unique GWAS-QTL overlaps separately for eQTLs, sQTLs, and caQTLs, and we report these in Table 1.

To test for overall enrichment of QTL overlapping with GWAS catalog SNPs, we downloaded the 1000 genomes VCF files (ftp://ftp.1000genomes.ebi.ac.uk/vol1/ftp/release/20130502/) and subsetted these to the EUR samples. We used vcftools to identify all SNPs in LD R^2^ > 0.8 with a GWAS catalog SNP and removed duplicate SNPs. We used our IPSDSN eQTL lead SNPs as input to SNPsnap (https://data.broadinstitute.org/mpg/snpsnap/), and computed 1000 random sets of SNPs using default parameters to match for LD partners, MAF, gene density, and distance to nearest gene. We determined the number of occurrences of eQTL lead SNPs in the GWAS catalog SNP + LD partners, and did the same for the 1000 matched SNP sets. The IPSDSN eQTL lead SNPs had more overlaps (92) than any of the matched sets (median: 58, range 37-87). Note that this number of overlaps is fewer than the number we report in Supplementary Table 11; this is because we detect more overlaps when using LD from our own samples than when using 1000 genomes LD patterns, which is expected since 1000 genomes EUR LD does not perfectly reflect LD in our data. We performed the same overlapping process for lead eQTL SNPs from each GTEx tissue, and plotted the number of overlaps per tissue in Supplementary Figure 21.

